# T7 phage-assisted evolution of riboswitches using error-prone replication and dual selection

**DOI:** 10.1101/2023.01.11.523572

**Authors:** Eduardo Goicoechea, Alfonso Jaramillo

**Affiliations:** Warwick Integrative Synthetic Biology Centre and School of Life Sciences, University of Warwick, Coventry CV4 7AL, UK; De novo Synthetic Biology Lab, i2sysbio, CSIC-University of Valencia. Parc Científic Universitat de València. Calle Catedrático Agustín Escardino, 9. 46980 Paterna. Spain

**Keywords:** Riboswitch, Enrichment, Genetic selection, Directed Evolution, Phage-assisted evolution

## Abstract

Leveraging riboswitches, non-coding mRNA fragments pivotal to gene regulation, poses a challenge in effectively selecting and enriching these functional genetic sensors, which can toggle between ON and OFF states in response to their cognate inducers. Here, we show our engineered phage T7, enabling the evolution of a theophylline riboswitch. We have replaced T7’s DNA polymerase with a transcription factor controlled by a theophylline riboswitch and have created two types of host environments to propagate the engineered phage. Both types host an error-prone T7 DNA polymerase regulated by a T7 promoter along with another critical gene - either cmk or pifA, depending on the host type. The cmk gene is necessary for T7 replication and is used in the first host type for selection in the riboswitch’s ON state. Conversely, the second host type incorporates the pifA gene, leading to abortive T7 infections and used for selection in the riboswitch’s OFF state. This dual-selection system, termed T7AE, was then applied to a library of 65,536 engineered T7 phages, each carrying randomized riboswitch variants. Through successive passage in both host types with and without theophylline, we observed an enrichment of phages encoding functional riboswitches that conferred a fitness advantage to the phage in both hosts. The T7AE technique thereby opens new pathways for the evolution and advancement of gene switches, including non-coding RNA-based switches, setting the stage for significant strides in synthetic biology.

## Introduction

Phages have transformed the directed evolution of genetic systems with their robust tolerance for high mutagenesis loads and swift replication rates. Phage Assisted Continuous Evolution (PACE) technology (1), leveraging the M13 phage, fast-tracks protein evolution in continuous cultures. This method substitutes an essential gene in the M13 phage genome with an inactively transcribed version in the host, allowing any gene of interest (GOI) to evolve if it can trigger the transcription of the critical gene. The PACEmid (2) technology, a subsequent refinement of the PACE approach, moves almost all phage genes to the host, stopping these genes’ adaptation. PACEmid aids the evolution of satellite phages by packaging plasmids as functional virions, or phagemids, using the phage’s packaging signal, proving particularly beneficial for GOIs with minimal initial activity. For example, it evolved the cro repressor into a transcription factor activator, creating the smallest known transcription factor activator of just 63 amino acids. PACEmid also increases the non-specific mutation rate tenfold. However, the use of M13 phage in bioreactors poses challenges. Post-infection, bacterial hosts survive, releasing non-evolved phages from biofilm-producing cells, a problem that intensifies under strict selection conditions. This becomes especially troublesome when the initial GOI activity is low or the GOI exhibits significant activity changes depending on the chemical environment, as is the case with gene switches.

We present T7AE, a new genetic selection system that utilizes phage T7 for the phage-assisted evolution of gene switches, notably riboswitches. Departing from prior reliance on M13 phage, we use the lytic T7 phage to spur the evolution of gene switches composed of non-coding RNA fragments. While earlier research primarily focused on evolving protein-based systems, non-coding RNA switches have seen significant development through computational design methods (3–6). Our study addresses this disparity, expanding phage-assisted evolution techniques to include gene switches. This extension provides a powerful experimental platform for the directed evolution of gene switches. Through combining phage-assisted evolution, predominantly applied to proteins, with the distinctive attributes of the lytic phage T7, we seek to uncover the evolutionary potential of gene switches.

Riboswitches, specific regions within mRNA molecules, possess a unique tertiary structure that impacts downstream gene expression (7). Binding to small molecules triggers a structural rearrangement in riboswitches, leading to modifications in the secondary structure of the succeeding mRNA sequence, either forming a new transcriptional terminator or revealing previously hidden ribosome binding sites (RBS) (8). Although riboswitches frequently occur in archaea and prokaryotes, they’ve been noted in eukaryotes and even prophages, as well as engineered to operate in viruses (9,10). Their cross-species functionality and significant mutability make riboswitches compelling research subjects, offering versatile applications and opportunities for generating novel variants that no longer recognize original ligands (11). Additionally, riboswitches’ compact size and clear operational mechanism make them one of the simplest genetic switches, illuminating principles of gene regulation. Investigation into riboswitches’ characteristics and applications holds immense promise for advancing synthetic biology and genetic engineering. Researchers have strived to uncover new riboswitches (12), enhance their functionality (13), and even design them *De novo* (14) due to their potential. Current methods for creating novel riboswitches include computer modeling (15), directed evolution (16), and *De novo* design (17). While many approaches concentrate on *In vitro* selection, requiring subsequent *In vivo* validation, there are instances of *In vivo* methods (18), which allow for the screening of viable riboswitch candidates early on, broadening the sequence space and amplifying the riboswitch’s final activity. The development of diverse and modular selection circuits is key to overcoming these challenges (19,20).

In this study, we focus on the accelerated evolution of genetic switches using dual selection methods, a critical process for effectively selecting both active and inactive genes in environments that induce ON and OFF states (21). Our primary objective is to engineer new riboswitches using lytic phages as the evolutionary platform. The theophylline riboswitch, our gene of interest, gets incorporated into the T7 phage genome and subsequently randomized using phage genome engineering techniques (22). This work represents the first instance of a functional riboswitch integration into a phage genome, setting it apart from prior studies identifying ribozymes in phages (23). We aim to evolve these riboswitches through T7AE and analyze the resulting phage population. The report is divided into four primary sections: an introduction to T7AE, detailing our dual selection method for directed evolution via T7 phage; a description of our riboswitch phage library; a presentation of our dual selection cycles; and finally, an estimation of riboswitch activity from the phage phenotype which demonstrates riboswitch adaptation during evolution. We follow these sections with a detailed discussion. In our experimental process, a varied phage population carrying randomized theophylline riboswitches undergoes several cycles of dual selection, leading to a noticeable reduction in riboswitch diversity within the phage population. This process also results in the emergence of new riboswitches with notable activation changes between selection stages. Our assessment of riboswitch sequence variability and measurement of phage activity under different inducer conditions offers insights into the evolutionary dynamics of riboswitches within the phage population and their functional responses to inducers.

## Results

### T7AE: Dual selection method for directed evolution via T7 phage

We engineer the T7 phage genome and its *Escherichia coli* host to establish an error-prone phage replication mechanism for implementing our T7AE system. We remove the DNA polymerase gene (gp5) on the T7 phage genome and replace it with a cI transcription factor from the λ phage, regulated by a theophylline riboswitch at its 5’ UTR. On the host side, we use an error-prone variant of gp5 (24) to enhance the T7 phage mutation rate. We also introduce a gene for the selection or counter-selection of functional or non-functional riboswitches, respectively. Our approach of combining an error-prone phage replication system with positive and negative selection steps is designed to encourage the emergence of novel, functionally enhanced riboswitch variants. This strategy, aimed at broadening our understanding of non-coding RNA engineering, allows for the experimental evolution of riboswitches. We outline our T7 phage-based screening strategy for novel riboswitch sequences in Figure 1. Utilizing homologous recombination (22,25) (Fig 1A), we integrate our DNA fragment of interest into the T7 phage genome. We create plasmids containing phage genome-homologous regions, aiming to remove the phage’s gp5 gene. The plasmids house either a positive control riboswitch sequence (PAJ341), or a library of riboswitch variants (PAJ342), as depicted in Figure S4 (further details in Supplementary Materials). The recombined gene fragment, carrying multiple elements (Fig 1B), includes a T7 promoter, a theophylline riboswitch version (26) (Fig. 1C), the λ phage-derived cI transcription factor (27), and the trxA gene, essential for phage replication. The trxA serves as a cloning marker to identify recombinant T7 genomes (22). During selection, the λ cI repressor activates the PRM promoter (28), which controls downstream gene expression. This configuration integrates the theophylline riboswitch-controlled gene expression system into the T7 phage genome, enabling the study and evolution of riboswitches within the framework of phage-assisted evolution.

**Figure 1.**
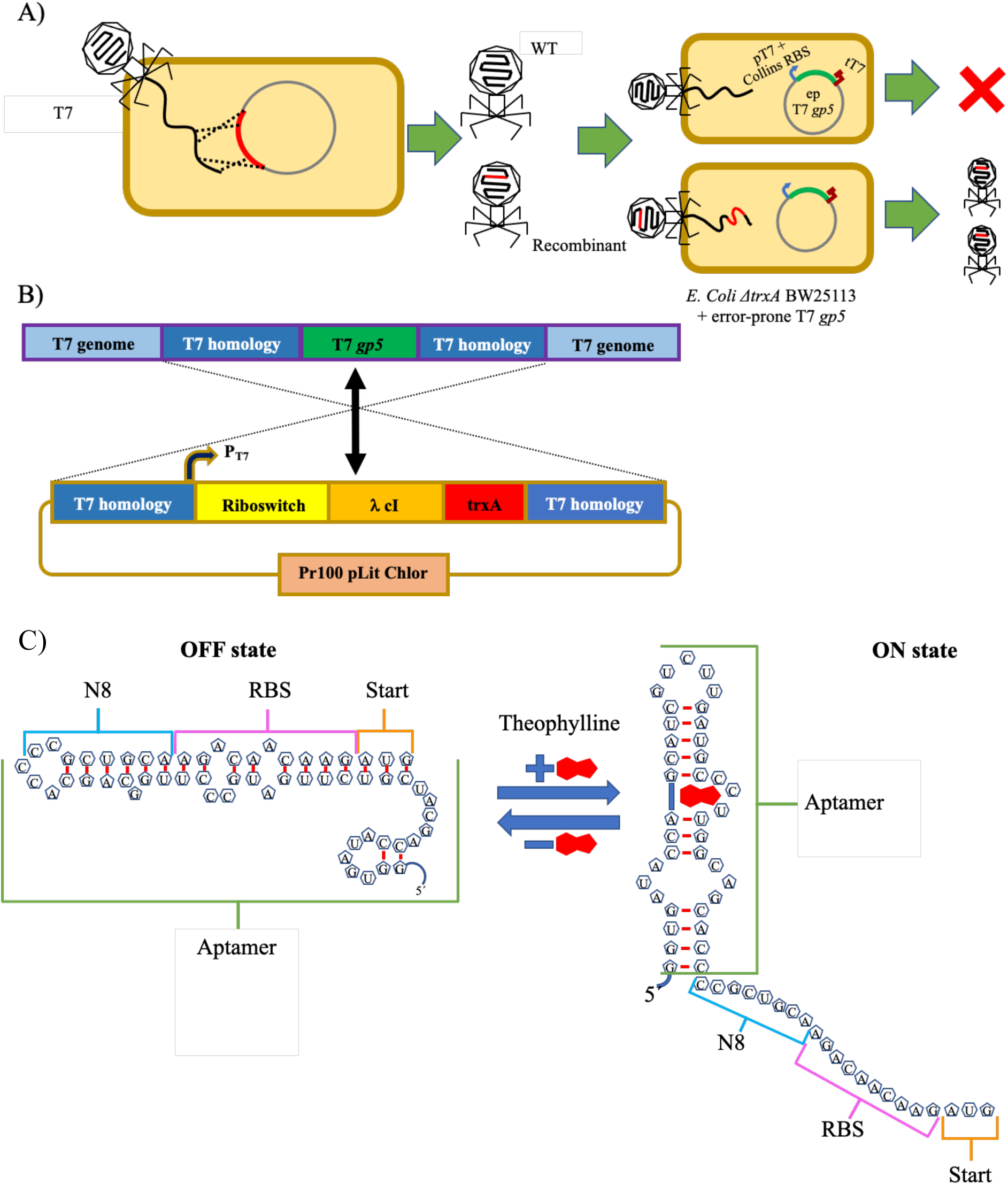
Engineering a riboswitch in the T7 phage using trxA marker. A) Obtaining recombinant phages: A schematic representation of the homologous recombination process occurring within the phage population. Recombinant phages are generated as a result, and these recombinants are selected by infecting a specific replication strain. B) Detail of genetic constructs: Depiction of the T7 genome and the recombination plasmid, highlighting the homology regions and the elements they flank. Homologous recombination enables the swapping of these two fragments. C) Versions of the theophylline riboswitch: Illustration of the “OFF” state (left) and the “ON” state (right) of the theophylline riboswitch.

To facilitate the evolution of a conditional genetic device, we employed a dual selection strategy consisting of positive and negative stages (29,30) (Fig. 2). For the positive selection phase, we added a high concentration of theophylline during phage replication, with the riboswitch controlling the replication rate via λ cI, thereby determining the selection outcome. We infected an engineered *Escherichia coli* strain, primed for positive selection, with our T7 phage carrying the riboswitch. This strain, carrying a cmk gene knockout essential for T7 phage replication (24) but not *Escherichia coli* growth, was transformed with a plasmid encoding the PRM promoter regulating the cmk gene. Theophylline’s presence selectively favored the phages carrying an active riboswitch, as these were the only ones capable of producing the λ cI necessary to activate the essential cmk gene.

**Figure 2.**
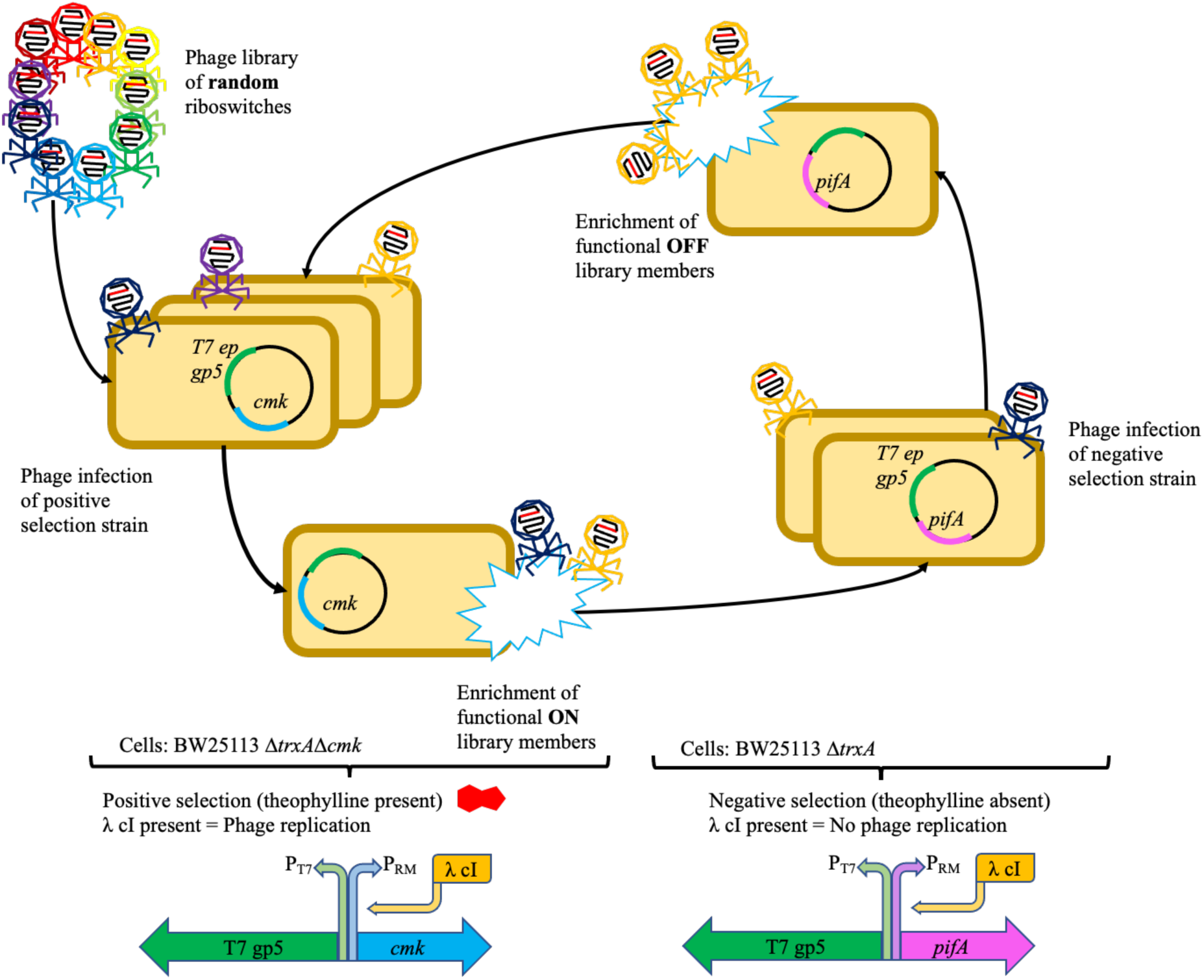
Schema showing the design and use of our T7AE system. General structure of the full phage evolutionary procedure: 1) Phage library infection: A phage library comprising various riboswitch variants infects a positive selection strain containing a positive selection plasmid. 2) Positive selection step: In the presence of theophylline, only phages carrying functional variants of the riboswitch will be able to replicate effectively. 3) Negative selection step: The surviving phages from the positive selection are subjected to a negative selection step. In the absence of theophylline, the riboswitches should remain inactive. However, to eliminate constitutively active phages, the pifA gene is transcribed, rendering these phages unable to replicate effectively. 4) Cycling between positive and negative selections: The resulting phages from the negative selection step infect the positive selection strain once again, initiating another cycle. This iterative process progressively exerts evolutionary pressure, driving the selection and improvement of functional riboswitch variants over time. The T7AE system combines positive and negative selection steps to enforce evolutionary pressure, facilitating the development and enhancement of riboswitches within the phage population

In the negative selection phase, a different *Escherichia coli* strain, retaining the original cmk gene and transformed with a plasmid containing the PRM-regulated pifA gene, was used for infection. The pifA gene, originating from the F-plasmid, inhibits T7 replication (31). This stage requires an absence of theophylline for the riboswitch to remain inactive. Any remaining riboswitch activity would escalate λ cI translation, thus activating the pifA gene and resulting in abortive infection. Consequently, phages carrying a riboswitch mutation that doesn’t fully repress λ cI translation in theophylline’s absence struggle to replicate properly. By alternating between positive and negative selection in theophylline’s presence and absence respectively, we progressively screened the initial pool of randomly mutated riboswitches (65,536 or 4^8^ variants) to yield more efficient riboswitches. This strategy fosters the enrichment and evolution of riboswitches with enhanced conditional behavior across selection cycles.

### Library design

To foster a broad library of phages, we infused random mutations into an 8-nucleotide region of the riboswitch, chosen based on previous research (26) (Fig. 3). The theoretical diversity of the library, considering all possible combinations of mutations in this 8-nucleotide region, encompasses 65,536 distinct variants. Given our transformation efficiency—measured at 1.11 * 10^6^ colony-forming units (cfu) per microgram (µg)—we accomplished a 17-fold coverage of the library with the utilized cultures (Fig. 3). This coverage implies a representation of the majority of library variants, thereby boosting the probability of unearthing novel and enhanced riboswitches through the evolutionary process.

**Figure 3.**
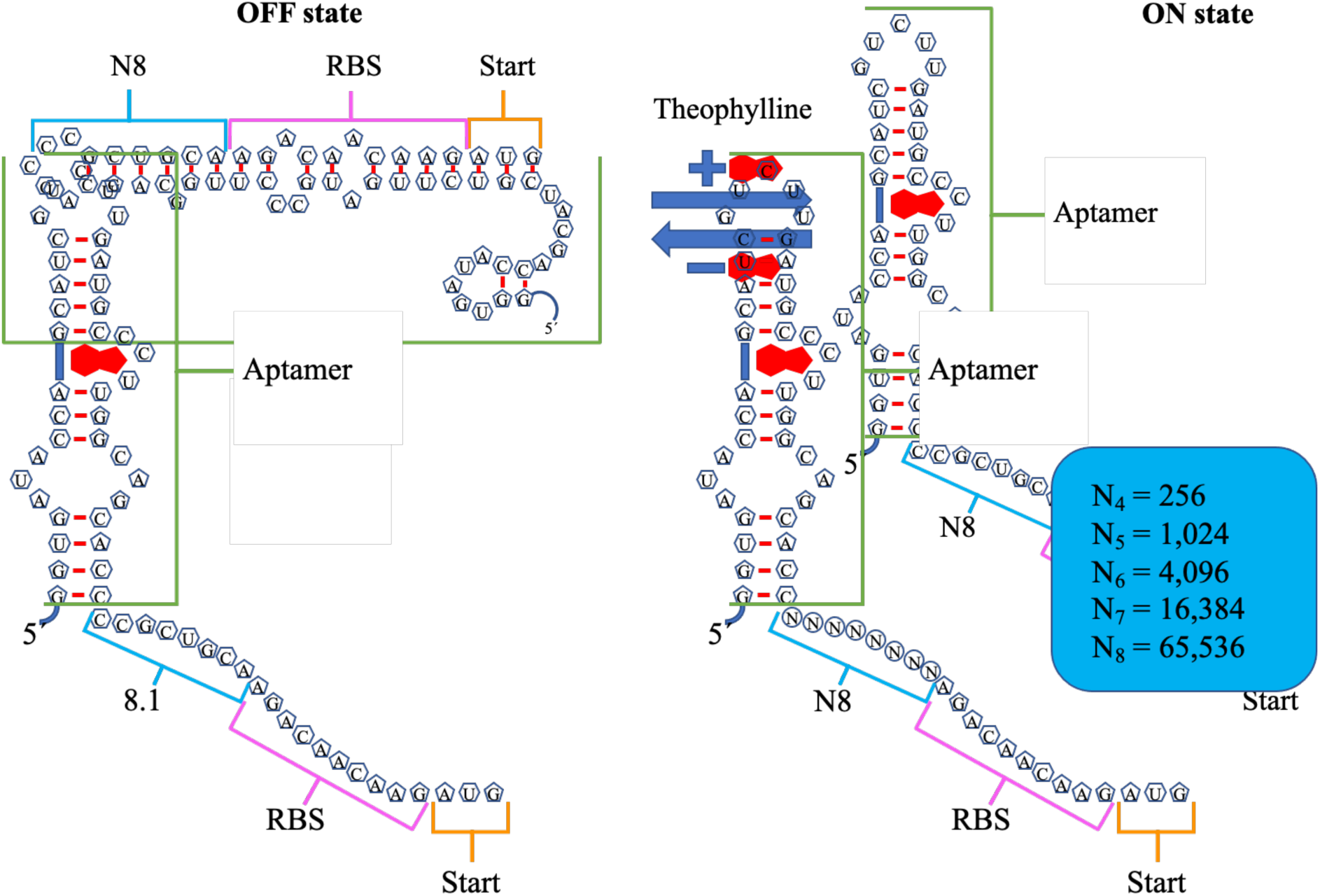
Design of riboswitch library. “OFF” state of the riboswitch (top left), “ON” state of the riboswitch (top right), 8.1 version of the riboswitch, which showed a 36-fold activation in a previous report(28) (bottom left) and structure of the theoretical library obtained after randomizing the 8 nucleotides following the stem of the riboswitch (bottom right).

To examine the initial diversity of the riboswitch libraries, we conducted sequencing analyses on samples taken from the various plasmid constructs and the phages post the homologous recombination process. Sequencing outcomes disclosed the existence of unique versions of the riboswitches in both the plasmid libraries (PAJ341 and PAJ342) and the phage library (Fig. 4). Significantly, no duplicate sequences were discerned in either the plasmid or the phage populations containing the randomized libraries. This attests that our method successfully yielded diverse and non-repetitive riboswitch variants, thereby establishing a robust groundwork for the subsequent selection and evolution process. Figure 4 offers a visual illustration of the sequence diversity observed in the plasmid and phage libraries.

**Figure 4.**
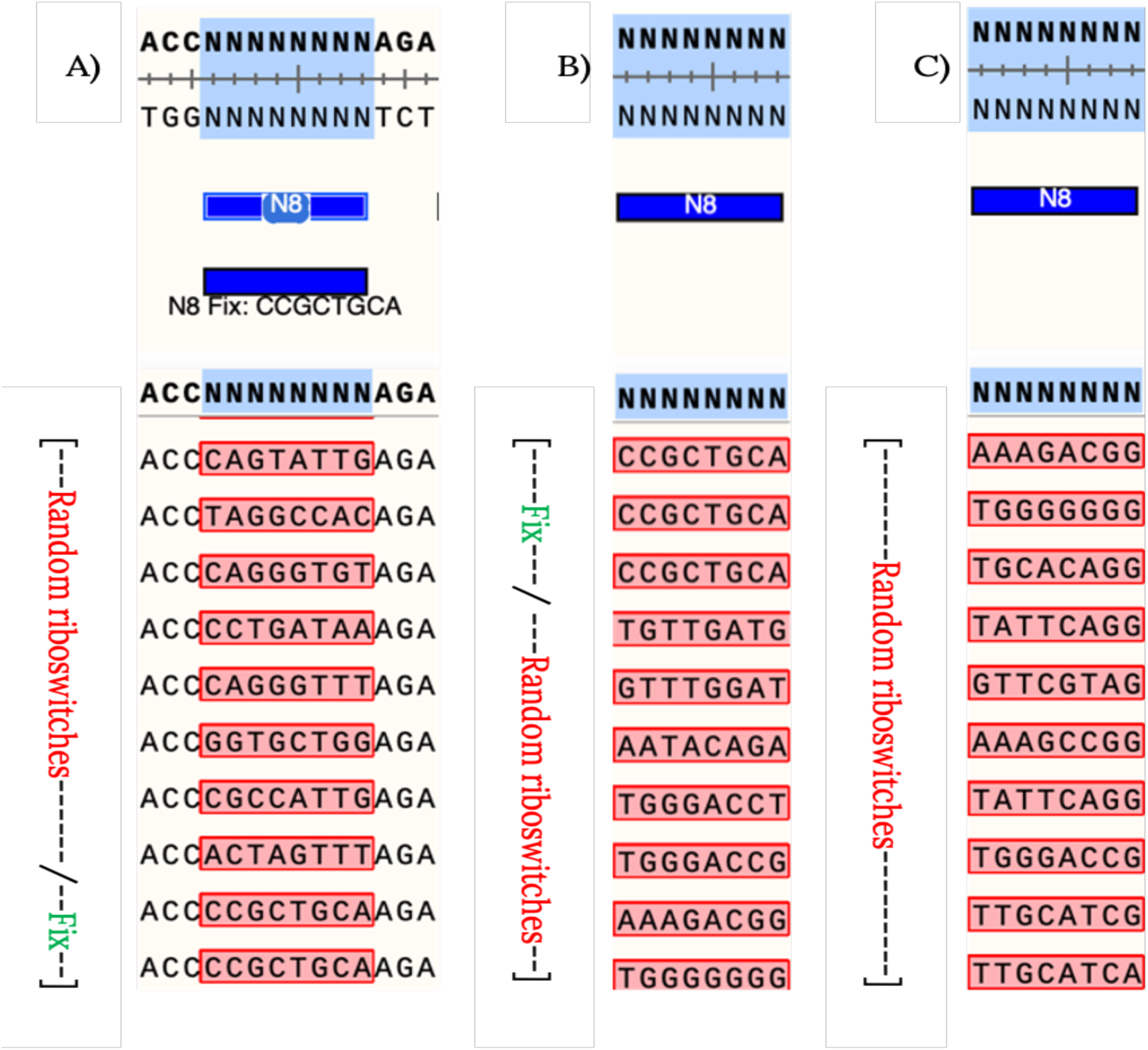
Sequences obtained from initial plasmid and phage libraries. Collection of riboswitch sequences obtained during the development of the different libraries. The sequences were obtained through Sanger sequencing of PCR products derived from single colonies carrying plasmids PAJ341 (positive control riboswitch) and PAJ342 (random library of switches). The riboswitch sequence used as a positive control (marked as “Fix” in the figure) served as a reference during the initial setup of the procedure. The sequences obtained from the plasmid libraries demonstrate the diversity of riboswitch variants generated, laying the foundation for subsequent selection and evolution steps.

### Dual selection cycles show enrichment with an engineered T7-phage library of randomized riboswitches

After carrying out 16 cycles of positive and negative selections, we analyzed a total of 24 libraries to assess the sequence variation and evolution of the riboswitches. During this analysis, we studied smaller library samples comprising 105 reads across the 16 selection cycles. This sample size was deemed appropriate as it encompassed the maximal diversity of the riboswitch library, containing 65,536 variants. The selection process entailed using 1.5 mM theophylline for the phage populations before the 10th generation, while 1 mM theophylline was employed for each generation thereafter to impose a more rigorous selection.

Figure 5 illustrates the outcome of the analysis, displaying diagrams of the 8 randomized nucleotides, encased by 5 fixed nucleotides on either side. The prominence of each nucleotide in the diagram directly reflects its overrepresentation at that position. This thorough analysis unfolded across multiple libraries, as enumerated in Table S1, and spanned diverse phases of the evolutionary trajectory. Each step carries the label “cmk” or “pifA”, identifying its association with either positive or negative selection. In the wake of the evolutionary process, these diagrams afford valuable insights, underscoring the dynamic shifts in nucleotide preferences and mutations within the riboswitch sequences.

**Figure 5.**
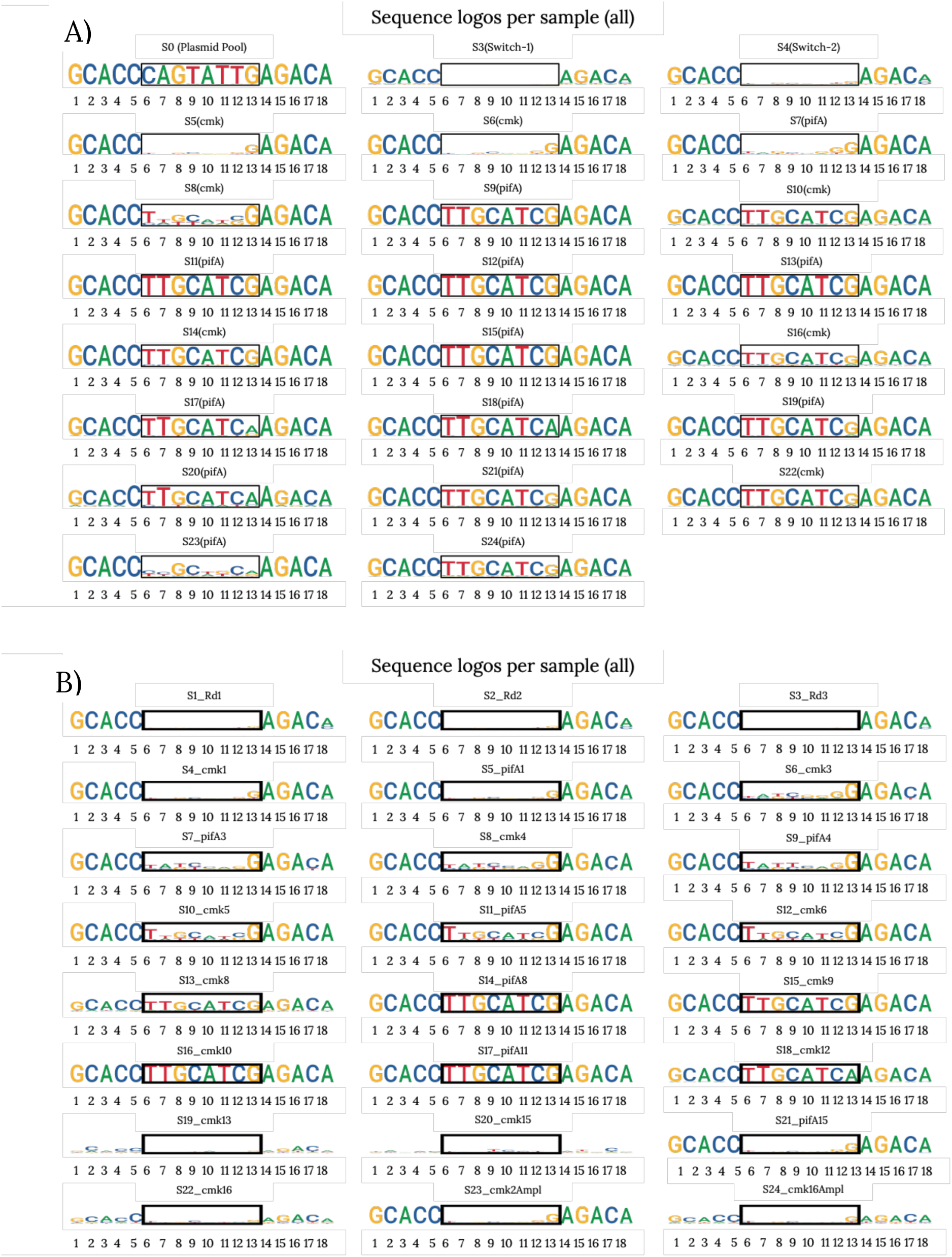
Evolution of predominant riboswitch sequences throughout dual selection cycles. Diagram representing the sequence variation for each library, arranged from the earliest library (top left) to the most recent library (bottom right). Each set corresponds to the predominant sequence observed in the phage population after each selection cycle. The black square indicates the region of the riboswitch sequence that was randomized. The size of each nucleotide in the diagram represents its relative abundance in the sample population. The nomenclature “S+number” indicates the generation number since the beginning of the project. Panel A of the figure shows the representation of the first set of libraries that were assessed. Initially, no specific sequence dominance was observed in the early selection cycles (Switch-1 and 2). However, over time, a progressive increase in the dominance of a particular sequence was observed, indicating its successful selection. Panel B of the figure represents a second set of libraries, focusing on the early generations to fill in the gaps observed in panel A and evaluate additional phage generations. Further details about the process can be found in the Materials and Methods section, and additional information regarding these experiments can be found in Table S1. Note that the last set of logos may have been affected by human error during preparation, which could explain the observed sequence variations.

Our next-generation sequencing (NGS) data analysis for various libraries yielded intriguing results. As demonstrated in Figure 5B, the initial set of sequences analyzed reveals no overt dominance of any sequence. However, a dwindling variation within each phage population becomes apparent with the advancement of selection steps, suggesting a convergent evolution towards a prevailing sequence. This pattern aligns well with past studies (9,28)that report a favored nucleotide, such as G, in position 8.

In a subsequent round of NGS scrutiny, most of the sequences echo this pattern of diminishing variation across generations alongside the rise of a dominant sequence. Yet, a subset of libraries (S23 in A and S19, S20, S21, S22, and S24 in B) display a heightened variation. The underpinnings of this amplified variation within these specific libraries currently remain elusive, warranting deeper exploration.

### Estimation of riboswitch activity from phage phenotype shows riboswitch adaptation during evolution

We executed a series of virulence index (VI) assays (32) to substantiate our selection results, as outlined in the Materials and Methods section. We tested a variety of phage populations procured throughout the process, such as the positive control (a recombinant phage bearing a known, functional riboswitch), phages from generations 2, 5, and 16, and a wild type T7 phage population lacking a riboswitch (negative control). Each population underwent testing under theophylline’s presence and absence, employing positive and negative selection strains. Our assays examined OD600 absorbance curves and regional virulence values, enabling us to calculate the virulence index. Figure 6 showcases a sample set of results derived from infecting the positive selection strain with phages subjected to 16 selection cycles..

**Figure 6.**
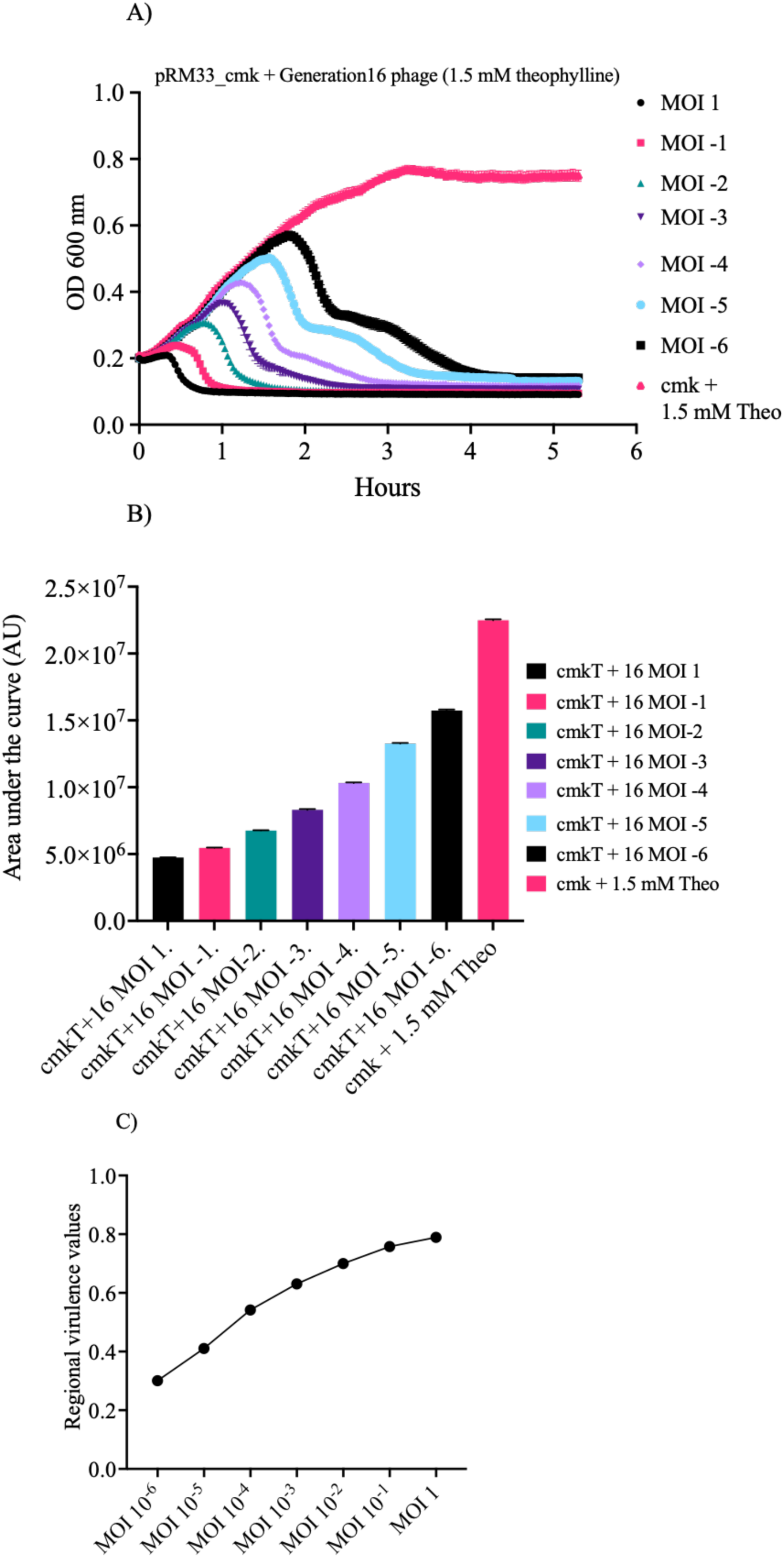
Standard set of data used to obtain the virulence index for a phage population infecting a specific strain. A) Killing curves of cell cultures with varying multiplicities of infection (MOIs), where different phages have been added, along with the growth curve of the cell strain without phages. The stationary phase is marked as the last timepoint included to calculate the area under the curves. B) Calculation of the area under each curve from panel A, represented as Arbitrary Units (AU) on the Y-axis. The cultures labeled as “cmkT” indicate the presence of theophylline in the media. C) Representation of the values obtained from panel B using Equation 1, as described in the Materials and Methods section. The value for the area under this curve corresponds to the virulence index for each phage in the respective strain.

We evaluated the activation efficiency of riboswitches across diverse phage populations, including wild-type T7, the Positive Control Riboswitch, and samples from cycles 2, 5, and 16. We used virulence index values from varying combinations of phages, strains, and theophylline concentrations to ascertain this efficiency. Our results presented unique patterns across these populations. While the wild-type T7 showed no change in its virulence index, the Positive Control Riboswitch generally indicated a higher fold-change in activation. Test samples from cycles 2, 5, and 16 revealed virulence variations. Particularly, in the negative selection step, phage populations from cycles 2 and 5 (selected at 1.5 mM) displayed higher activation rates than the Positive Control. The population from cycle 16 (selected at 1 mM) showed lower efficiency. However, when the riboswitch underwent conformational changes, cycle 16 at 1 mM manifested the most significant activation rate, showing a 28% increase. These observations underscore the variable activation efficiency of riboswitches in different phage populations and underscore the significant role of selection conditions in their performance.

Figure S1 (33) provides a comparative view of the potential impact of changes in riboswitch sequences on virulence through their predicted structures. It not only features random sequences from the initial selection stages and the Positive Control riboswitch sequence, but also the dominant sequences emerging in the selection’s later stages. This comparison serves to underscore the structural variances among the riboswitch variants obtained throughout the selection process and their possible influence on virulence.

## Discussion

Our novel genetic system, T7AE, successfully facilitated riboswitch selection through T7 phages and a dual-selection process in live cells, leading to the evolution of a theophylline riboswitch. The sequencing of the resulting phage populations offered a timeline of evolution, pinpointing when specific nucleotides solidified their positions and when a certain sequence rose to dominance in the population (Figure 5). From our findings and data in Table S1, it is evident that phages surviving the initial selection exhibited a preference for G as the 8th nucleotide in the sequence. By the sixth cycle, the TTGCATCG sequence (or its slightly improved version, TTGCATCA) emerged as the dominant variant, maintaining prevalence through the following generations. This dominant sequence bears notable similarities to the control sequence (CCGCTGCA) from earlier studies (26), with six of the eight purines and pyrimidines occupying identical positions.

We utilized virulence index calculations to evaluate each phage population’s infection efficiency, providing a robust comparative measure among diverse phage populations and a more thorough assessment than growth curves alone (32). Figure 6 showcases the virulence index values of various phage populations, including two controls: Wild Type T7 and the Control riboswitch (26) depicted in Figure 2B. We utilized Wild Type T7 to assess switch activation and its virulence impact, while the Control riboswitch served as a functional standard. Predictably, the Wild Type T7 virulence index hovered around 30% (Figure 6A and B), given the absence of an activatable switch. The positive control phages, used to establish the procedure, exhibited a virulence increase of 20% and 17% in the positive selection, while no difference surfaced in the negative selection.

Despite the observations in Figure 6 panels A and B, our statistical analysis found that differences between the riboswitch’s ON and OFF states weren’t always statistically significant. In the positive selection, regardless of the theophylline concentration, the positive control consistently exhibited significant differences between the ON and OFF states. The negative control, in alignment with expectations, showed no significant differences in any situation. Generation 5, among the various phage populations examined, presented no significant differences between the ON and OFF states. However, generations 2 and 16 showed significant differences in activation states (generation 2) and when comparing activation efficiencies across various theophylline concentrations (generations 2 and 16). These results demonstrate that riboswitch populations undergo adaptations over successive generations, leading to variations in activation efficiencies and virulence.

Our findings can be attributed to the incomplete understanding of the pifA molecular cascade (34,35)and the potential for system leakage. Given the current limited knowledge of the pifA protein mechanism and the phage exclusion process (34–36), some effects may go unnoticed during the selection process. This could result in more significant variations between results, especially in the negative selection..

Our examination of the actual test populations and their virulence index (VI) values revealed intriguing patterns when compared to the phage carrying the known control riboswitch sequence. Two of three test populations exceeded the VI values of the control riboswitch sequence. However, as Figure 7 demonstrates, this increased infectivity didn’t correlate with improved riboswitch sequences. Population 5, despite presenting the highest virulence values, didn’t exhibit improved and even showed reduced activation efficiency, particularly in the positive selection. This pattern could stem from the prevalence of a sequence perpetually in the “ON” state, sufficiently abundant to outperform other sequences during growth assays. No sequence in generation 5 achieved complete fixation within the population as the NGS analysis (Figure 5, S8 in A and S10 in B) reveals, hence the improvements in virulence indexes with additional selection steps. Our T7AE genetic system has proven its potential in evolving theophylline riboswitches. The combination of sequencing analysis of phage populations and VI calculations delivers rich insights into the phage population evolution timeline and infection efficiency. Nonetheless, statistical analyses show not all differences between the riboswitch’s ON and OFF states are statistically significant, implying the need for deeper exploration of the pifA molecular cascade and possible system leakage. The observed inconsistency between higher VI values and activation efficiency underscores the selection process’s complexity and the need for comprehensive characterization of riboswitch populations to identify the most effective sequences.

**Figure 7.**
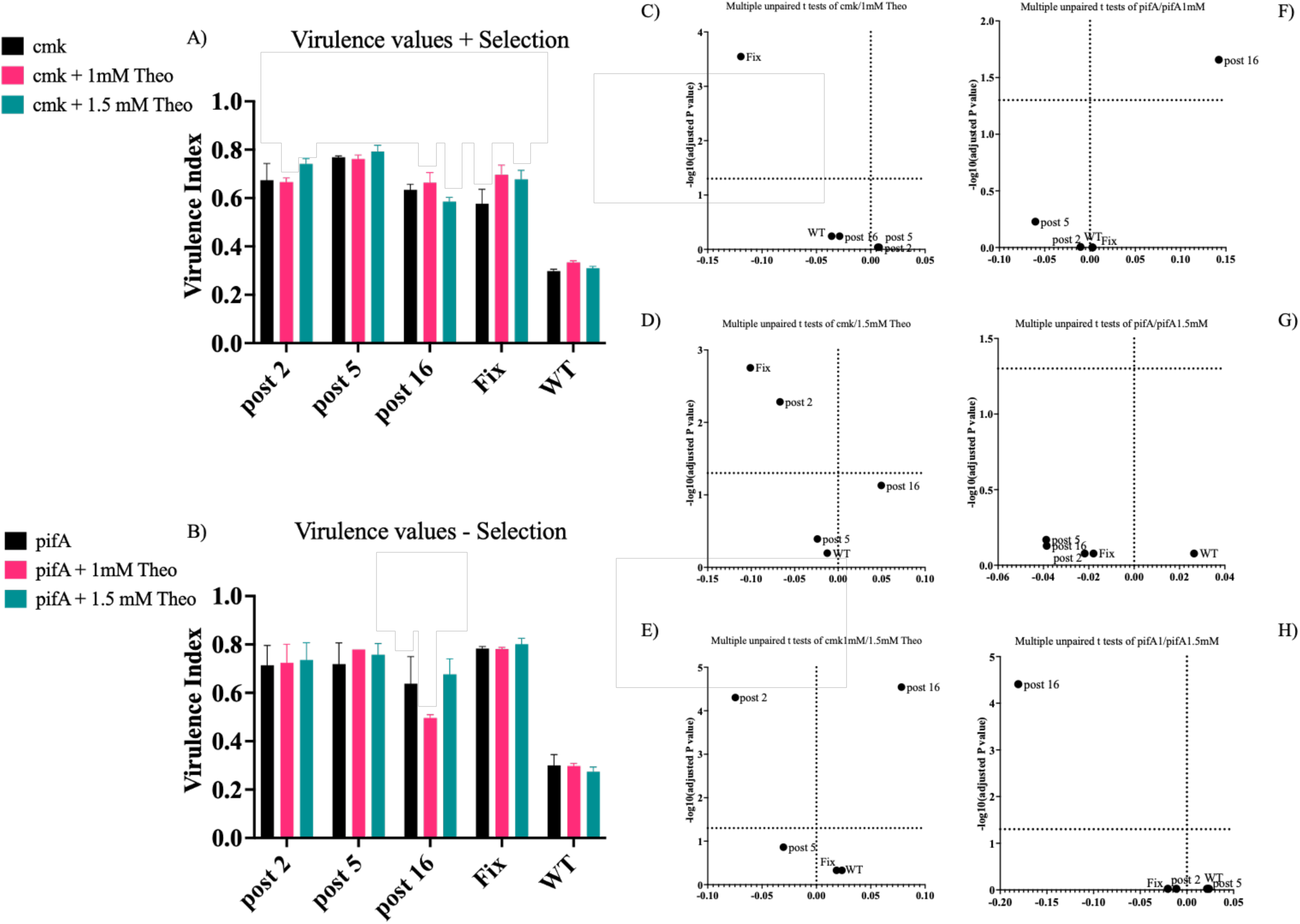
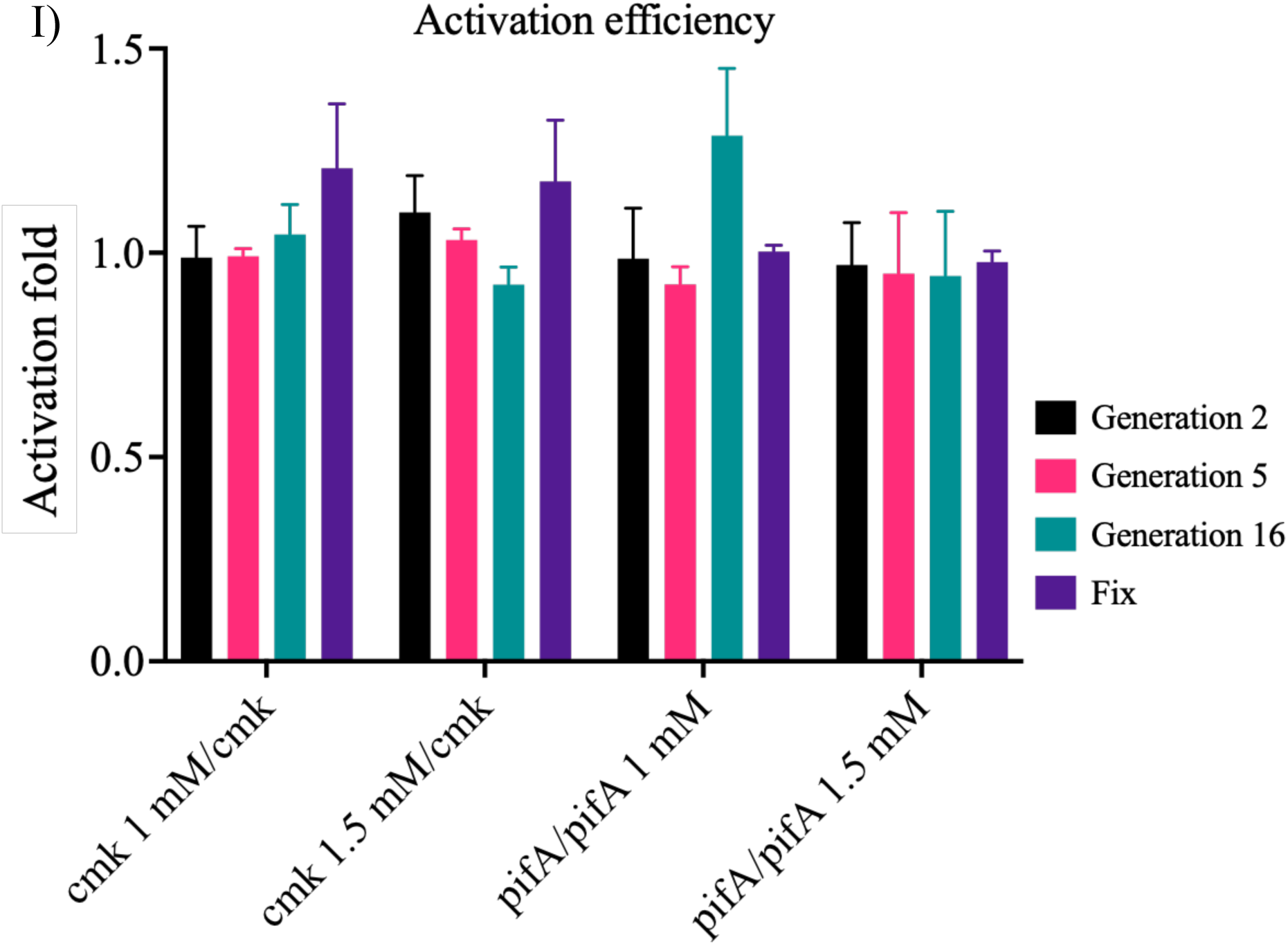
Virulence comparisons between phage populations. Results of the virulence index calculations for different phage populations (Wild Type/Negative control, Control riboswitch/Positive control, and after 2, 5, or 16 cycles of selection) when infecting the positive and negative selection strains in the presence and absence of different concentrations of theophylline. The virulence index values range between 0 and 1, with higher values indicating higher virulence of the phage population. The X-axis represents the presence or absence of theophylline in the cultures, denoted by a “T.” A) The virulence indexes of the five phage populations are shown after infecting the positive selection strain in the absence and presence of theophylline. In the presence of theophylline, the riboswitch changes to an “ON” conformation, leading to higher virulence. B) Virulence index for the different phage populations when infecting the negative selection strain in the absence and presence of theophylline. In this case, virulence is expected to be higher in the absence of theophylline as the riboswitch remains in the “OFF” state. When theophylline is added and the switch turns “ON,” the expression of pifA blocks efficient T7 replication, reducing virulence. C to H) Depiction of p-values representing the statistical significance of the pair comparisons between different conditions in both positive and negative selections. The corresponding statistics can be found in Table S4. Panel I shows the fold increase in virulence for the different concentrations of theophylline used in both positive and negative selections. In the positive selection, the virulence values in the presence of theophylline were divided by the values in its absence, while the opposite calculation was done for the negative selection. The significance of figures C-H was determined using unpaired t-tests, and the error bars in panel I were obtained using error propagation.

Our T7AE research marks a significant step forward in riboswitch evolution by incorporating a riboswitch into a T7 phage’s genome. Despite potential for further technical enhancements, we strongly believe in the potential impact of our approach on riboswitch and biosensor evolution. Our selection strategy’s efficacy in identifying effective riboswitch sequences from a random pool is strongly supported by our combined virulence index and next-generation sequencing (NGS) data analysis, as demonstrated in Figures 4 and 6. Interestingly, phage virulence does not necessarily correlate with riboswitch efficiency. Some phages demonstrated heightened virulence over subsequent generations despite seemingly less efficient riboswitch sequences. These outcomes may result from the presence of constitutive sequences that confer an advantage during specific selection steps or particular sequences affecting regulation unexpectedly. Importantly, the riboswitches we obtained exhibit improved characteristics compared to the control riboswitch validated by previous literature (26), as seen in Figures 6 and 7. In the future, our technology could adapt to the PACEmid approach, evolving satellite T7 phages. The T7 phage’s successful use in evolution studies (37) inspires confidence; only the evolving DNA needs the packaging signal. Our T7AE riboswitch selection approach allows us to discover riboswitches with unique ligand specificities. Our dual selection process establishes a versatile methodology for the phage-assisted accelerated evolution of gene switches. In short, our T7AE system represents an innovative approach for riboswitch evolution, with great potential to advance riboswitch and biosensor development. The integration of T7 phages and the use of virulence index and NGS data contribute valuable insights into the selection process and resulting riboswitch populations. With further developments and exploration, our T7AE strategy holds significant promise for the production of functional riboswitches and the acceleration of gene switch evolution.

## Materials and methods

### LB media

All bacterial strains were grown in LB media for the different experiments, with any alternative media used only for specific experiments, and described accordingly. The formulation for 1L includes 10 g Peptone 140, 5 g yeast extract, and 5 g sodium chloride, diluted in ddH2O.

### Saline-Magnesium buffer

Solution used to store phage samples, avoiding the risk of contamination by microorganisms (38). For 1L, 50 mM Tris-HCl 7.5, 100 mM NaCl, 100 mM MgSO_4_ diluted in ddH20.

### Cell strains

Cells from the Keio Collection (39), were used to transform the confirmed plasmids and to use the resulting cells for the positive and negative selections. Strain specifications are described in the Supplementary Materials.

### Plasmids

Plasmid specifications and assembly details are described in the Supplementary Materials.

### Primers and gBlocks

All gene sequence constructs produced by Integrated DNA Technologies and can be found in Table S2. Melting temperatures (T_m_) of the primers were calculated using the NEB calculator. PCR reactions were carried out using Phusion Master Mix from NEB in a final volume of 25 µL for diagnostic PCRs, and 50 µL for purification PCRs.

### Digestion enzymes

All used enzymes were part of the FastDigest line from Thermo Scientific. All digestions were done at 37 °C, using 200 ng of DNA template, 1 µl of FastDigest enzymes, and 10x buffer solutions from ThermoFisher in a final volume of 20 µL.

### Polymerase Chain Reaction

All PCR reactions used for plasmid assembly were done using the 2x Phusion mix from NEB, in a 50 µL volume, according to manufacturer specifications. In cases where the secondary structure of the sequences interfered with the amplification, different percentages of DMSO were used (3, 6 %) to help the correct amplification. Diagnostic PCRs from colonies or phage samples were done using a 2x Taq or GreenTaq Mix (Thermo Scientific) under manufacturer’s specifications.

### Goldengate assembly

Based on Engler, Kandzia and Marillonnet (40). The desired plasmids were amplified via PCR. The primers used contain an overhang not present in the original template, which introduces a BsaI/Eco31I cutting site. Compatible ends were included into the tails of primers used to amplify the desired fragment introduced in the backbone. Using 100 pmol of each DNA template, 1 µL of BsaI (New England Biolabs/NEB), 1 µL of T4 DNA (Fermentas), 2 µL of 10x ligase buffer and H2O up to 20 µL, each reaction tube underwent the following series of cycles: 2 minutes at 37°C followed by 5 minutes at 16°C, 35 cycles, 5 minutes at 50°C and finally 5 minutes at 80**°**C.

### Agarose electrophoresis

Amplified samples were run in 1% agarose gels (41), to confirm their size and approximate quantity, at 95V for 30 minutes for most experiments. Smaller samples were run in 2% agarose gels.

### Purifications

All PCR reactions were purified using the Thermo GeneJET PCR purification kit. In cases where several bands appeared when running the sample in a gel, the GeneJET Gel extraction kit was used.

### Transformations

Plasmids were transformed into both electrically and chemically competent cells. For transformations in electrically competent bacteria, a minimum of 100 ng of DNA were added to 50 µL of competent cells and left on ice for 15 minutes, along with an electroporator cuvette. The volume was then transferred to said cuvette, the cells electroporated and then grown in LB media for 1 hour before being plated. For chemically competent cells, a minimum of 100 ng of DNA was added to the 50 µL of cells, which were left on ice for 30 minutes, then heat shocked at 42°C for 90 seconds, and returned to ice for 10 minutes before recovery in LB media for an hour at 37°C. Cells were then plated in an LB agar plate with their corresponding antibiotic resistances.

### Growth

To obtain cell cultures for the various assays, colonies picked from grown plates were grown overnight in 5 ml of LB with the appropriate antibiotic, using Falcon 15 ml round bottom tubes. In the cases of positive selection strains, to prepare them for the assays, 300 µL of 25 mM Theophylline were added to the 5 ml of LB. Overnight cultures were refreshed the next day and used when an OD between 0.2 and 0.3 was reached. The remaining overnight culture was used to create glycerol stocks.

### Phage killing and homologous recombination

To obtain phage populations or test their killing efficiencies, phages were added to cultures within the growth phase (OD 0.2-0.3), making sure the phages would be able to properly replicate. In cases where the objective was to produce homologous recombination, a rate of 1 phage per 10^5^ cells was used. These tests required phages to infect Δ*trxA* cells containing gp5/T7 DNA polymerase. This ensured that only recombinant phages would be able to replicate, as trxA is an essential protein for the process (24). All lysates were then centrifuged for 10 min at 3000 rpm and filtered with 2 µm filters, obtaining clean phage samples.

### Selection procedure

8 different strains were tested for each of the selection steps, using the control riboswitch to find the most efficient strain to be used for evolution. This was done by infecting each of the strains of interest with the recombined T7 phages in the following ways:

- <u>Growth assays</u>: In a 96-well plate, each well is filled with 180 µL of cells at an OD of 0.2-0.3, and 20 µL of the tested phages in a decreasing order along the length of the plate. In the case of cultures with theophylline, two ways were tested: 180 µL of a culture already containing theophylline, or 163.5 µL of culture plus 12.5 µL of 25 mM theophylline, to which the phages were added. As a way to keep all records the same, and have a consistent concentration in the cultures, the first method was preferred. The plate reader (model, make) took OD measurements every 2 minutes for a duration of 5 hours. To compare the differences between strains and select the best candidates for the evolution procedure, the growth curves for each cell strain were plotted using Microsoft Excel and Graphpad Prism 9.
- <u>Plaque assays</u>: These assays were developed to check the replication efficiency of the phages (42). By infecting 300 µL of cells at an OD of 0.2-0.3 with 100 µL of phage at different dilutions, mixing it with 3 ml of soft LB agar (LBA 50% in LB), and then plating it, clear spots or “plaques” can be observed in the bacterial lawn of the plate. A higher number of plaques indicates a higher number of phages, indicating which strains were better for the replication of the phages. These assays were made both in the presence and absence of 1.5 mM Theophylline, to assess the “leakiness” of the system and the activation of the riboswitch.
- <u>One-Step assay</u>: These assays (Figure S2) follow the same procedure as a plaque assay, but phage samples were obtained from infecting a culture (43). Samples were taken every 4 minutes over a period of 40 minutes, showing the rise in the number of phages as the infection progresses. The assays were also made in the presence or absence of 1.5 mM theophylline to determine the differences due not only to the two types of sample treatment, but to the addition of the activating molecule to the media, thus affecting the phages’ replication process. These phage samples were then used for plaque assays as described previously.

### Next-Generation Sequence Analysis

This strategy was designed following the 16S amplicon sequencing protocol from Illumina, using its Nextera XT DNA Library Kit (Illumina #FC-131- 001) to prepare the samples for sequencing in an Illumina MiSeq (Figure S2). A first PCR was carried out to amplify a 175 bp fragment from the phage genome containing the closest elements present upstream and downstream from the region containing the 8 randomized nucleotides of the riboswitch (Figure S3A). The primers used in this PCR also incorporated sequences necessary to carry out the indexing PCR for the sequencing (Figure S3B). Combinations of primers for the Index PCR are shown in Table S2.

### Riboswitch sequence analysis

Due to the lack of available computing power, the initial set of analyses were done in a fraction of the total number of sequences, approximately 10^5^ per sample.

Sequence quality control was performed using the FastQC software (44). After trimming, all reads corresponding to each sample were aligned to the initial “parent” sequence (as designed and cloned into the initial vector prior to selection) using a recursive Needleman-Wunsch pairwise alignment algorithm (45). Once aligned, all reads were trimmed from both ends until only the 8 nt-long riboswitch sequence plus 5 bases on either side remained. Trimming was performed using base R, by finding the 8 nucleotide long “NNNNNNNN” motif in the parent sequence and extracting this position (±5bp) for all reads in each sequence. Once extracted, nucleotide proportions were calculated by stacking each position and counting the number of each nucleotide that was informative (not ambiguous), which is considered a Bit. The program does not only count which percentage of each nucleotide is present in every position, as there are many reads showing an ambiguous read or just a gap in these positions. To account for this, it creates a list and only counts the number of times every base is clear, represented as an informative Bit, according to information theory. The entire process of nucleotide counting and plotting was performed using the R package *ggseqlogo* (46).

### Evolution assessment via virulence index calculation

The selected procedure shown previously (32,47), was used, by which a value known as the virulence index was ascribed to different phages to compare them to one another. The virulence index was obtained via a two-step process:

I. Calculating the local virulence (υ1) via the following formula:

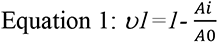

where A_0_ represents the area under the growth curve of a cell culture without phages until reaching stationary phase, and A_i_ the area under the curve of a culture at a certain MOI until the same timepoint. υ1 is measured in a range of 0 to 1, with 0 being no virulence whatsoever, and 1 being the maximum theoretical virulence, instantaneous death of the infected cells.
II. Once the local virulences over a range of MOIs were assessed, they were represented on a curve. The area under that curve (A_p_) was calculated following the formula

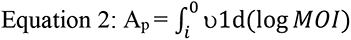

with *i* being the base10 log of the lowest MOI, which in this case was -6.
III. That value was then divided by the theoretical maximum value (A_max_) for the same curve, giving the virulence value for a specific phage on a specific strain. In this case, this maximum value corresponded to 6.
IV. These values were readily comparable with each other, and were taken both as they were, and to calculate the “activation efficiency” for the riboswitches inside the phages, indicating the fold increase between states of the switch in those cases.

The virulence index measurements were made for 5 different phage populations: 3 at different stages of selection, 1 for the original riboswitch-carrying phage, and one for WT T7. Each of these phages was used to infect the positive and negative selection strains under three conditions: 0 mM, 1 mM and 1.5 mM Theophylline; to assess their virulence under each condition. All calculations were made using Microsoft Excel 2019 and Graphpad Prism 9. Statistical significance between the different virulence indexes was done via unpaired t-tests.

### Riboswitch Activation metric

This is a value representing the activation fold of the riboswitch in each of the populations. This was obtained by dividing the virulence values for a particular strain in cases where the switch should be “ON”, by the value in the case where it should be “OFF”. This allows to see the fold increase in virulence between the presence/absence of theophylline and thus, the activation/inactivation fold of the riboswitch.

## Supporting information

The supporting information includes a list and description of used plasmids and their assembly methods, and a list of relevant primers and their sequences, as well as diagrams of several experimental procedures and nomenclatures.

## Abbreviations

*cmk*: CMP kinase
*trxA*: Thioredoxin A
MFE: Minimum free energy
MOI: Multiplicity of infection
PACE: Phage Assisted Continuous Evolution
SELEX: Systematic Evolution of Ligands by Exponential enrichment

## Acknowledgements

We thank WISB for the use of their facilities during this work and Prof. David Hodgson for a gift of the T7 phage. EG was supported by a Marie Sklodowska-Curie early training fellowship. This work was supported by the European Union’s Horizon 2020 research and innovation programme under ITN Marie Sklodowska-Curie (grant agreement 642738 [MetaRNA]), the Biotechnology and Biological Sciences Research Council (grant BB/P020615/1 [EVO- ENGINE], the Spanish Ministerio de Ciencia e Innovacióńs (grant PID2020-118436GB-I00 [GeneCircuits++]) and the European Innovation Council (grant 101070948 [PhotoSynH2]) to AJ.

## Supplementary Materials and Methods

### Strain List

Cells from the Keio Collection (1), namely the parental strain BW25113 (SAJ128), and KO for *trxA* (SAJ130) and both genes *cmk* and *trxA* (SAJ19) (CGSC, http://cgsc2.biology.yale.edu). The Keio Collection cells were used to transform the confirmed plasmids and to use the resulting cells for the positive and negative selections.

### Plasmid List

Below is a list of the different plasmids utilized throughout the project, with their official denomination, their name throughout the project, and the serial number in the lab’s records. Their specific assembly is discussed in the next section.

- Pr100 pLit Chlor/EG_001/PAJ343: 3494 bp, with a pMB1 origin of replication and Chloramphenicol resistance. Genes 4.7 and 5.3 from the T7 phage’s genome; both flanking a T7 promoter followed by the gene for thioredoxin A to incorporate them via homologous recombination.
- pSEVA631/EG_002/PAJ86: 3001 bp, containing the default multiple cloning site flanked by an rrnB T1 terminator and a lambda t0 terminator, a pBBR1 origin of replication, and a Gentamicin resistance. This plasmid was used as the backbone for all selection plasmids. Parental positive selection plasmid (EG002+): 7135 bp Parental negative selection plasmid (EG002-): 8677 bp
- HRF plasmid/EG_cI01/PAJ341: 3569 bp, based upon the Δ*cmk* version of Pr100 pLit Chlor. Via the use of GoldenGate assembly, a gene fragment containing a T7 terminator, a T7 promoter, and a fixed version of the theophylline riboswitch with a high-fold activation, taken from (2), as seen in Figure 1B. This plasmid was used as a positive control for the efficiency of the method.
- HRR plasmid/EG_cI02/PAJ342: 3569 bp. Exactly the same elements as EG_c01, but with the riboswitch contains a randomised region of 8 nucleotides after the stem of the riboswitch and before the RBS (Figure 1C). Thus, a plasmid library of riboswitches containing theoretically 65,536 variants. These will be used for experiments once the method has been properly set up for the fixed version of the riboswitch.
- Positive selection variants: Based upon the positive selection plasmid, different combinations of promoter and RBS were tested to select for the ideal one to use in the current experimental design. This meant a combination of promoter and RBS that gave very low production of *cmk* for a riboswitch in an “OFF” state (without theophylline), and a high one for an “ON” state riboswitch (with theophylline). 8 variants were used, 6 of them test strains, and 2 of them used as controls.

`o` The positive control/EG_cI03/SAJ782 (6093 bp) contained a T7 promoter and a strong synthetic RBS in control of the *cmk* gene, ensuring its expression and the proper replication of T7 phages, regardless of the presence of theophylline.
`o` The negative control/EG_cI10/SAJ789 (6054 bp) contained neither promoter nor RBS in front of *cmk,* so phages would not be able to replicate under any circumstances, as the transcription system could not be initiated in this case
`o` The test variants/EG_cI04-09/SAJ783-8 (6155 bp) contained the previously described pRM promoter followed by 1 of 6 different RBSs of different strengths. These RBSs were known as B0030, B0031, B0032, B0033, B0064, and B00Syn (2).
- Negative selection variants: As in the positive selection, 8 variants were tested, 2 of them being controls and 6 of them being tests. All the variations are as previously described, except for the plasmids containing *pifA* instead of *cmk*.

`o` Positive control/EG_cI11/SAJ700: 7635 bp.
`o` Negative control/EG_cI18/SAJ707: 7600 bp.
`o` Test variants/EG_cI12-17/SAJ701-6: 7700 bp.

### Plasmid assembly specifications

All the constructs assembled were based on previously available plasmids present in the laboratory’s stock. The assemblies were made either by enzymatic digestion at 37 °C and overnight ligation at 20°C, or using the process known as GoldenGate ligation.

- PAJ343: The Homologous Recombination plasmid was based on a pre-existing plasmid present in the lab, Pr100 pLit Chlor, itself based on pLITMUS 28 (https://www.addgene.org/vector-database/1528/); and containing the genes for CMP/dCMP kinase and thioredoxin A, making it 4226 bp. The plasmid was modified further before assembling the final version, eliminating the *cmk* gene via enzymatic digestion, making it 3490 bp.
- PAJ86: Based upon the European standard of SEVA plasmids (3), allowing for easy to reproduce modularity. This original plasmid underwent further modifications to assemble the positive and negative selection plasmids. Such modifications included the addition of an error-prone version of T7 phage’s DNA polymerase controlled by a T7 promoter in the 5’ direction, and BsaI sites flanking an R0010 promoter and Ribosome Binding Site (From here onwards RBS) controlling RFP in the 3’ direction, followed by B0010 and B0012 terminators; followed by the selection gene (*cmk* or *pifA* for the positive and negative selections, respectively).
- Selection plasmids: All different cargos were introduced into EG002 via Goldengate assembly, using the previously mentioned GoldenGate cutting sites

---

**Figure S1.**
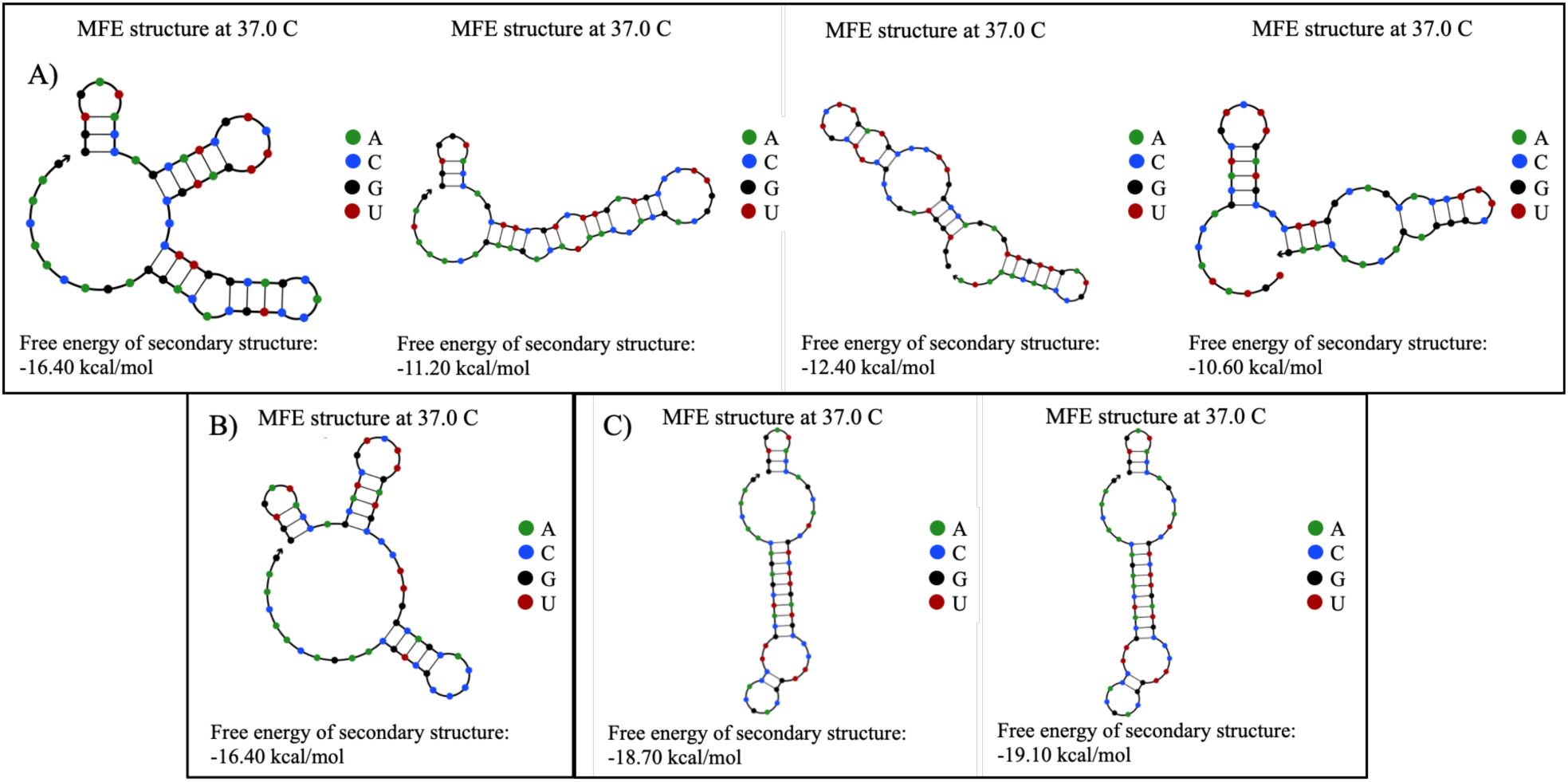
Structural comparison between riboswitch sequences. Using NUPACK (4), the secondary structures corresponding to different sequences throughout the selection process were predicted, as well as their MFE (minimum free energy). **A)** Sequences from the random pool of riboswitches from the early stages in the selection. **B)** Riboswitch used as a standard and positive control, with a fixed sequence (CCGCUGCA). **C)** Sequences that were overly represented by the final steps of the selection, as seen in Figure 3B. Left structure: TTGCATCG, Right structure: TTGCATCA. This shows that starting from a random pool of switches, we have been able to obtain sequences which managed to rival and even surpass the stability of the positive control switch.

**Figure S2.**
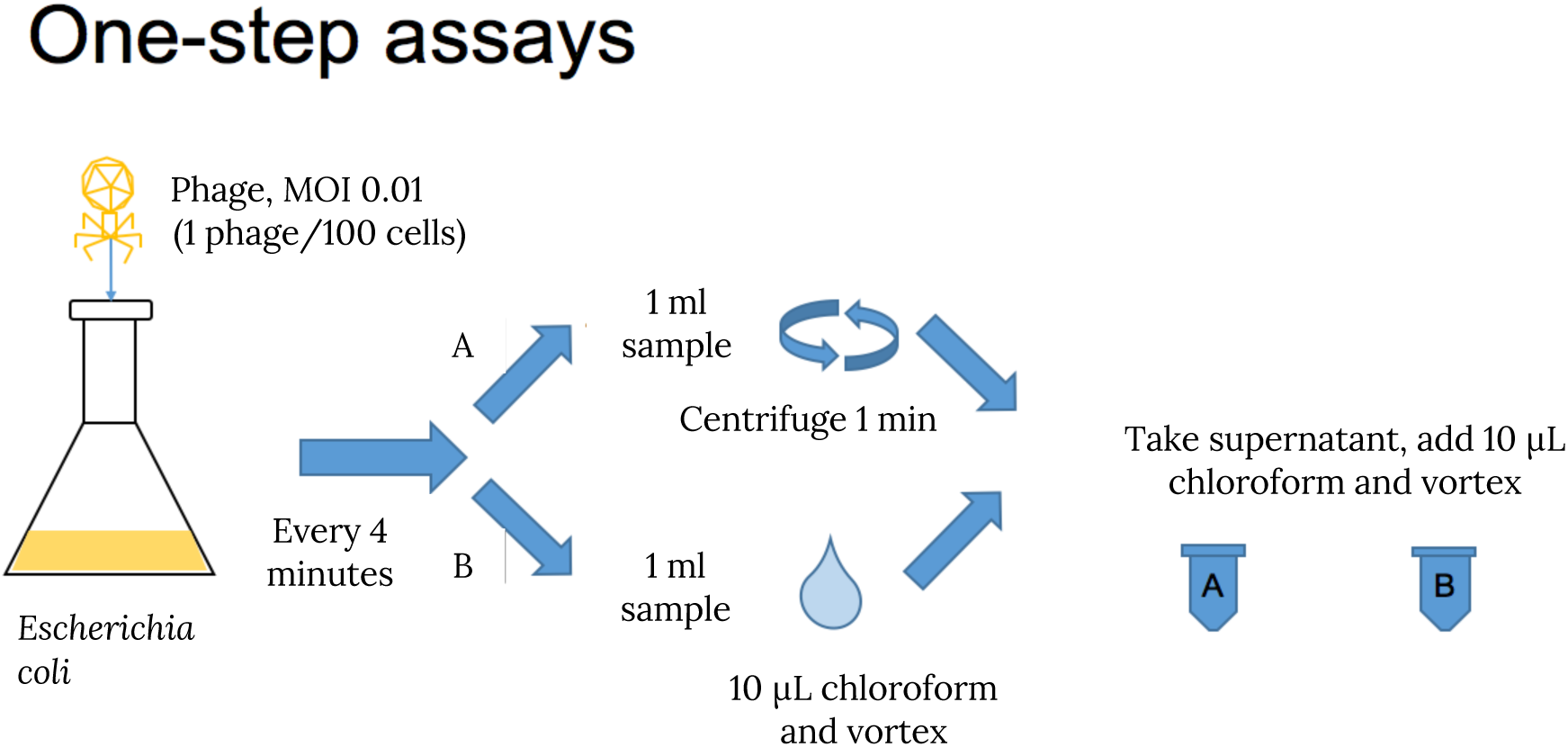
General diagram of a one-step assay. Once the phages, at a MOI (Multiplicity of Infection) of 10^-2^ (1 phage/100 cells) have been added to the culture, samples are taken and undergo different treatments. “A” are centrifuged to pellet the cells and obtain the phages present in the supernatant, while “B” samples have Chloroform added to them, causing bacteria to burst and release phages that had not yet exited the cells. To eliminate any remaining bacteria present in the final samples, Chloroform is added again, and those phage samples are then used for a plaque assay, where a difference should be observable between different samples and time points.

**Figure S3.**
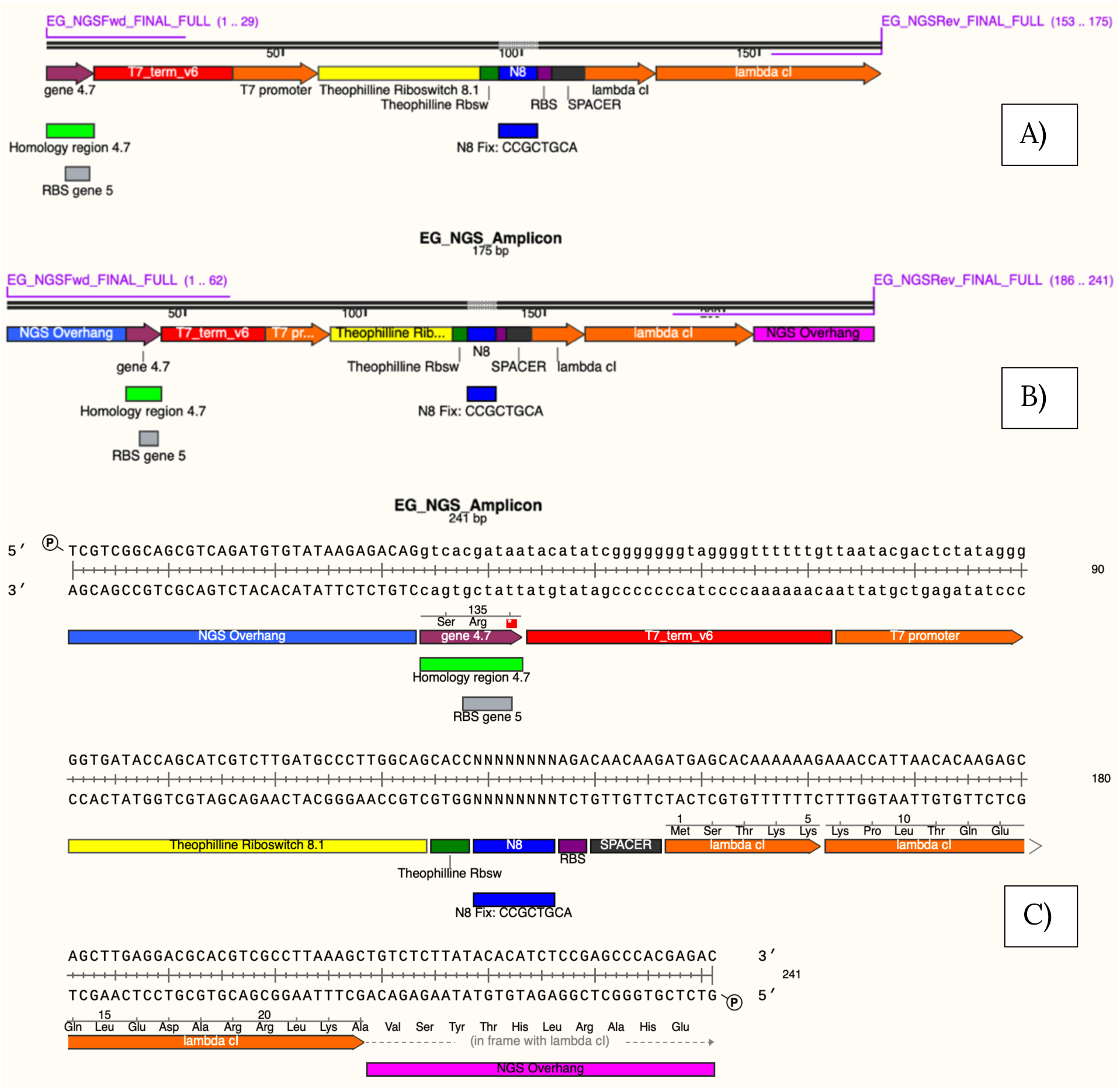
Amplification steps for NGS preparation. Selected amplicon to be used for the NGS procedures. A) Original amplicon as it is present in the recombined phages’ genome, 175 base pairs in size, containing fragments upstream and downstream from the random riboswitch sequence. B) Amplicon obtained after the PCR, with the desired primers having added the required Illumina overhangs for the Indexing PCR. C) Detailed sequence of the final amplicon, indicating the original sequences and the overhangs.

**Figure S4.**
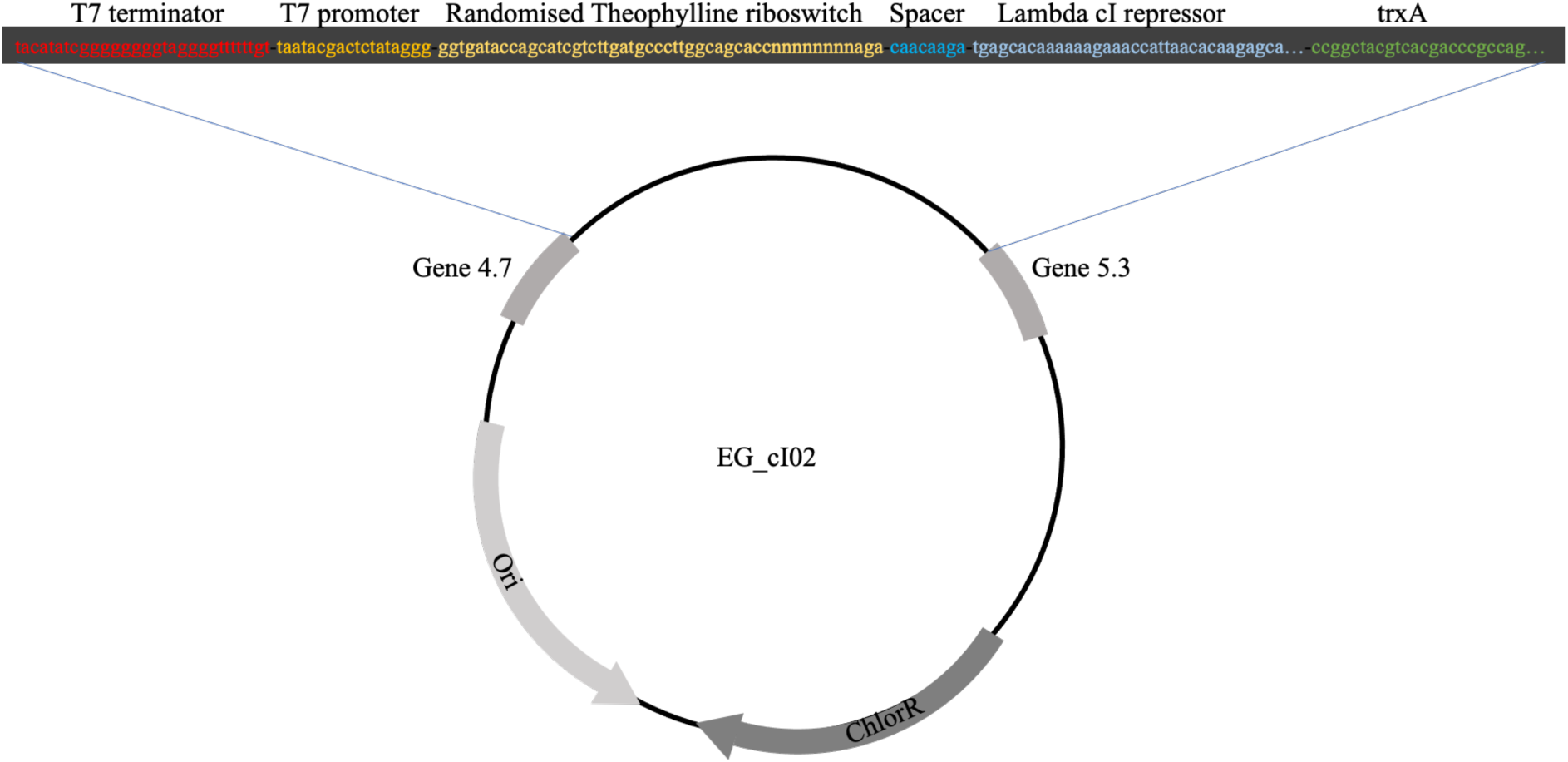
Schematic representation of HR plasmid EG_cI02. This is the version of the homology plasmid that was used for the creation of the library, carrying the fragment of interest flanked by two T7 genes to act as homology regions.

**Table S1.**
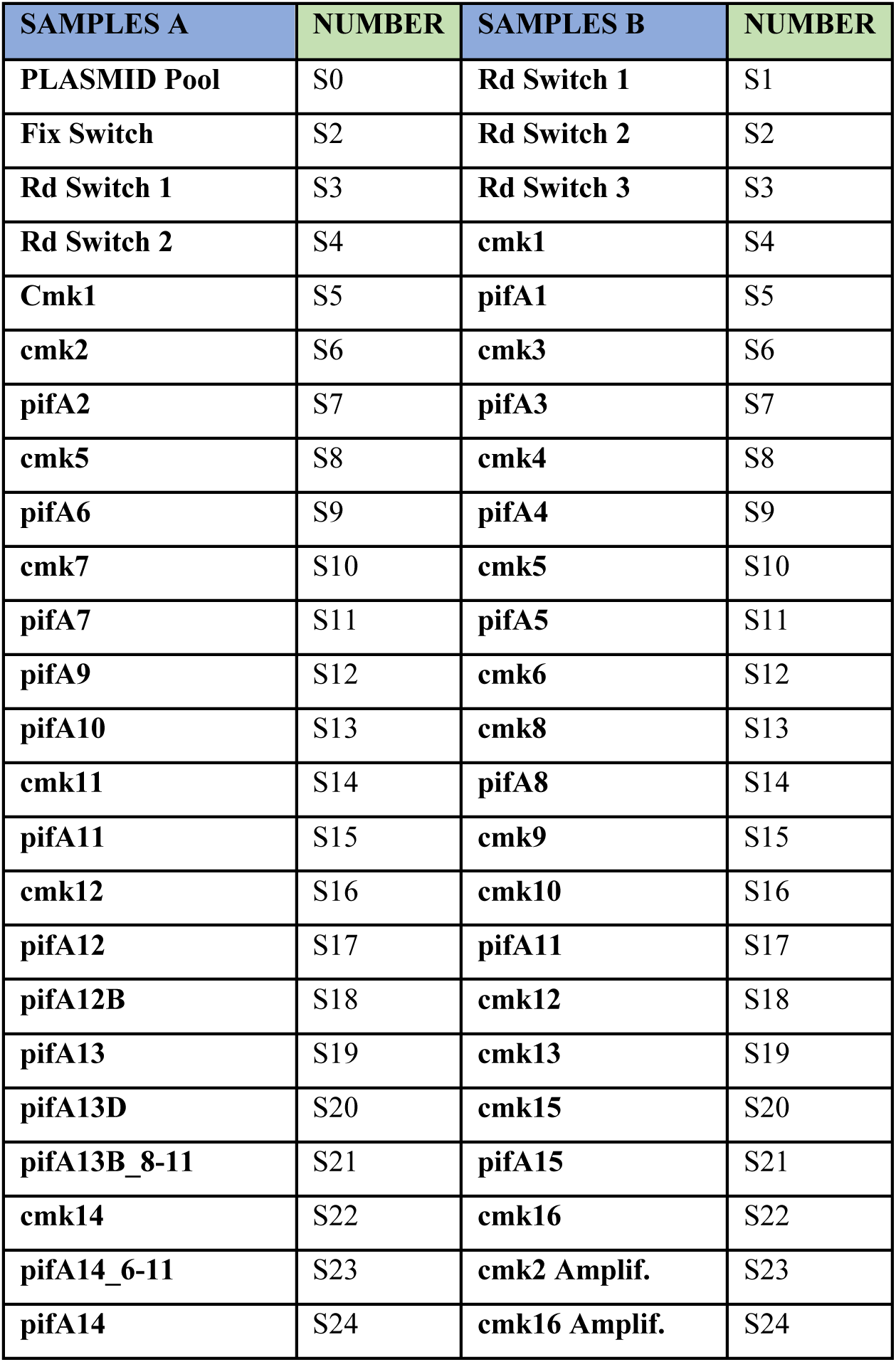
NGS sample nomenclature. Correspondence between the generation number of the different analysed libraries and the nomenclature used in the sequence variation diagram. cmk or pifA indicated whether the sample corresponded to a positive or negative step in the selection, and the number indicated the specific generation.

| **SAMPLES A** | **NUMBER** | **SAMPLES B** | **NUMBER** |
| --- | --- | --- | --- |
| **PLASMID Pool** | S0 | **Rd Switch 1** | S1 |
| **Fix Switch** | S2 | **Rd Switch 2** | S2 |
| **Rd Switch 1** | S3 | **Rd Switch 3** | S3 |
| **Rd Switch 2** | S4 | **cmk1** | S4 |
| **Cmk1** | S5 | **pifA1** | S5 |
| **cmk2** | S6 | **cmk3** | S6 |
| **pifA2** | S7 | **pifA3** | S7 |
| **cmk5** | S8 | **cmk4** | S8 |
| **pifA6** | S9 | **pifA4** | S9 |
| **cmk7** | S10 | **cmk5** | S10 |
| **pifA7** | S11 | **pifA5** | S11 |
| **pifA9** | S12 | **cmk6** | S12 |
| **pifA10** | S13 | **cmk8** | S13 |
| **cmk11** | S14 | **pifA8** | S14 |
| **pifA11** | S15 | **cmk9** | S15 |
| **cmk12** | S16 | **cmk10** | S16 |
| **pifA12** | S17 | **pifA11** | S17 |
| **pifA12B** | S18 | **cmk12** | S18 |
| **pifA13** | S19 | **cmk13** | S19 |
| **pifA13D** | S20 | **cmk15** | S20 |
| **pifA13B_8-11** | S21 | **pifA15** | S21 |
| **cmk14** | S22 | **cmk16** | S22 |
| **pifA14_6-11** | S23 | **cmk2 Amplif.** | S23 |
| **pifA14** | S24 | **cmk16 Amplif.** | S24 |

**Table S2.** Primer combinations for each NGS sample. Matrix indicating which primers correspond to each specific phage library in each of the two repetitions. The 6x4 matrix gives the possibility of separating between 24 samples at once, each of them with their own specific signal, thanks to the specific indexes present in each of the 2 primers used for the amplification. The different denominations for each library are based on the selection step they were obtained from, followed by the number of the generation. If there are multiple libraries corresponding to the same generation, a letter or date is included after the denomination. E.g. Sample cmk11 corresponds to the phage library obtained after the 11^th^ positive selection step in the evolutionary process. A and B correspond to the 1^st^ and 2^nd^ set of sequenced libraries, respectively.

| A) | N701 | N702 | N703 | N704 | N705 | N706 |
| --- | --- | --- | --- | --- | --- | --- |
| S502 | PLASMID<br>Pool | Fix<br>Riboswitch | Rd Switch 1 | Rd Switch<br>2 | Cmk1 | cmk2 |
| S503 | pifA2 | cmk5 | pifA6 | cmk7 | pifA7 | pifA9 |
| S504 | pifA10 | cmk11 | pifA11 | cmk12 | pifA12 | pifA12B |
| S517 | pifA13 | pifA13D | pifA13B_8-11 | cmk14 | pifA14_6-11 | pifA14 |
| B) | N701 | N702 | N703 | N704 | N705 | N706 |
| S502 | Rd1 | Rd2 | Rd3 | Cmk1 | pifA1 | cmk3 |
| S503 | pifA3 | cmk4 | pifA4 | cmk5 | pifA5 | cmk6 |
| S504 | cmk8 | pifA8 | cmk9 | cmk10 | pifA11 | cmk12 |
| S517 | cmk13 | cmk15 | pifA15 | cmk 16 | cmk2 Ampl | cmk16 Ampl |

**Table S4.** Full statistical data for virulence index significance. Explanation of the type of statistical analysis and corrections made to determine whether the differences between different populations were significant. **A)** Positive selection. Showing the pair comparison between all different populations: cells grown in the absence of theophylline (cmk), cells grown in 1mM theophylline (cmk + 1mM Theo), and cells grown in 1.5 mM theophylline (cmk + 1.5mM Theo). **B)** Negative selection. Showing the pair comparison between all different populations: cells grown in the absence of theophylline (pifA), cells grown in 1mM theophylline (pifA + 1mM Theo) and cells grown in 1.5 mM theophylline (pifA + 1.5 mM Theo). All calculations were done with Graphpad Prism 9.

|  |  |  |  |  |  |  |  |  |  |
| --- | --- | --- | --- | --- | --- | --- | --- | --- | --- |
| Test details |  |  |  |  |  |  |  |  |  |
| Test name | Unpaired t test |  |  |  |  |  |  |  |  |
| Variance assumption | Single pooled variance |  |  |  |  |  |  |  |  |
| Multiple comparisons | Set P value threshold |  |  |  |  |  |  |  |  |
| Method | Holm-Šidák method |  |  |  |  |  |  |  |  |
| Alpha | 0,05 |  |  |  |  |  |  |  |  |
| Number of tests performed | 5 |  |  |  |  |  |  |  |  |
| Number of rows omitted | 0 |  |  |  |  |  |  |  |  |
| **A)** | **Below threshold?** | **P value** | **Mean of cmk** | **Mean of cmk +**<br>**1mM Theo** | **Difference** | **SE of difference** | **t ratio** | **df** | **Adjusted P Value** |
| post 2 | No | 0,705422 | 0,6738 | 0,6661 | 0,007694 | 0,02024 | 0,3802 | 50 | 0,913224 |
| post 5 | No | 0,751494 | 0,7684 | 0,762 | 0,006444 | 0,02024 | 0,3184 | 50 | 0,913224 |
| post 16 | No | 0,201953 | 0,6344 | 0,6631 | -0,02867 | 0,02217 | 1,293 | 50 | 0,571508 |
| Fix | Yes | 0,000057 | 0,5767 | 0,6962 | -0,1195 | 0,02715 | 4,401 | 50 | 0,000283 |
| WT | No | 0,190931 | 0,2978 | 0,3338 | -0,036 | 0,02715 | 1,326 | 50 | 0,571508 |
|  | **Below threshold?** | **P value** | **Mean of cmk** | **Mean of cmk +**<br>**1.5mM Theo** | **Difference** | **SE of difference** | **t ratio** | **df** | **Adjusted P Value** |
| post 2 | Yes | 0,001303 | 0,6738 | 0,7407 | -0,06689 | 0,01963 | 3,407 | 50 | 0,005202 |
| post 5 | No | 0,229343 | 0,7684 | 0,7923 | -0,02389 | 0,01963 | 1,217 | 50 | 0,406087 |
| post 16 | No | 0,025309 | 0,6344 | 0,5848 | 0,04958 | 0,0215 | 2,306 | 50 | 0,074021 |
| Fix | Yes | 0,000353 | 0,5767 | 0,6777 | -0,101 | 0,02634 | 3,835 | 50 | 0,001763 |
| WT | No | 0,637125 | 0,2978 | 0,3103 | -0,0125 | 0,02634 | 0,4746 | 50 | 0,637125 |
|  | **Below threshold?** | **P value** | **Mean of cmk +**<br>**1mM Theo** | **Mean of cmk +**<br>**1.5mM Theo** | **Difference** | **SE of difference** | **t ratio** | **df** | **Adjusted P Value** |
| post 2 | Yes | 0,000012 | 0,6661 | 0,7407 | -0,07458 | 0,01485 | 5,022 | 38 | 0,00005 |
| post 5 | No | 0,048098 | 0,762 | 0,7923 | -0,03033 | 0,01485 | 2,042 | 38 | 0,137466 |
| post 16 | Yes | 0,000006 | 0,6631 | 0,5848 | 0,07825 | 0,01485 | 5,269 | 38 | 0,000029 |
| Fix | No | 0,38397 | 0,6962 | 0,6777 | 0,0185 | 0,021 | 0,8808 | 38 | 0,467438 |
| WT | No | 0,270232 | 0,3338 | 0,3103 | 0,0235 | 0,021 | 1,119 | 38 | 0,467438 |

| **B)** | Below threshold? | P value | Mean of pifA | Mean of pifA + 1mM Theo | Difference | SE of difference | t ratio | df | Adjusted P Value |
| --- | --- | --- | --- | --- | --- | --- | --- | --- | --- |
| post 2 | No | 0,765117 | 0,714 | 0,7244 | -0,01046 | 0,03487 | 0,3001 | 62 | 0,987041 |
| post 5 | No | 0,200169 | 0,7188 | 0,7792 | -0,06033 | 0,04659 | 1,295 | 62 | 0,590746 |
| post 16 | Yes | 0,004439 | 0,6379 | 0,4958 | 0,1421 | 0,04812 | 2,953 | 62 | 0,022 |
| Fix | No | 0,957819 | 0,7831 | 0,7805 | 0,002556 | 0,04812 | 0,05311 | 62 | 0,997813 |
| WT | No | 0,953239 | 0,3003 | 0,2975 | 0,002833 | 0,04812 | 0,05888 | 62 | 0,997813 |
|  | Below threshold? | P value | Mean of pifA | Mean of pifA + 1.5 mM Theo | Difference | SE of difference | t ratio | df | Adjusted P Value |
| post 2 | No | 0,453006 | 0,714 | 0,7357 | -0,02179 | 0,0289 | 0,7541 | 80 | 0,836338 |
| post 5 | No | 0,203151 | 0,7188 | 0,7576 | -0,03878 | 0,03022 | 1,283 | 80 | 0,678723 |
| post 16 | No | 0,289006 | 0,6379 | 0,6765 | -0,03856 | 0,03612 | 1,067 | 80 | 0,744457 |
| Fix | No | 0,620705 | 0,7831 | 0,801 | -0,01794 | 0,03612 | 0,4968 | 80 | 0,836338 |
| WT | No | 0,466713 | 0,3003 | 0,2739 | 0,02642 | 0,03612 | 0,7313 | 80 | 0,836338 |
|  | Below threshold? | P value | Mean of pifA + 1mM Theo | Mean of pifA + 1.5 mM Theo | Difference | SE of difference | t ratio | df | Adjusted P Value |
| post 2 | No | 0,672187 | 0,7244 | 0,7357 | -0,01133 | 0,02659 | 0,426 | 44 | 0,943301 |
| post 5 | No | 0,524939 | 0,7792 | 0,7576 | 0,02156 | 0,03364 | 0,6409 | 44 | 0,943301 |
| post 16 | Yes | 0,000008 | 0,4958 | 0,6765 | -0,1807 | 0,03568 | 5,064 | 44 | 0,000039 |
| Fix | No | 0,568476 | 0,7805 | 0,801 | -0,0205 | 0,03568 | 0,5746 | 44 | 0,943301 |
| WT | No | 0,51203 | 0,2975 | 0,2739 | 0,02358 | 0,03568 | 0,661 | 44 | 0,943301 |

